# Effect size of delayed freezing, diurnal variation, and hindgut location on the mouse fecal microbiome relative to a standardized biological variable

**DOI:** 10.1101/2023.07.13.548851

**Authors:** K.L. Gustafson, Z.L. McAdams, A.L. Russell, R.A. Dorfmeyer, G.M. Turner, A.C. Ericsson

## Abstract

**Background:** While murine fecal collection is central to microbiome research, there are a number of practical considerations that may vary during fecal sample collection, including time to sample storage, time of day the sample is collected, and position within the colon during terminal collections. While the need to control these factors is recognized, the relative effect on microbial community of duration at room temperature, time of day, and hindgut position, in the context of a known biological variable, is unclear. To answer these questions, and assess reproducibility of results across different microbiome compositions, parallel experiments were performed to investigate the effect of those factors on the microbiome of age- and sex-matched isogenic mice colonized with two different vendor-origin microbiomes.

**Results:** 16S rRNA amplicon sequencing data from flash-frozen fecal samples showed no statistical difference in alpha or beta diversity compared to samples incubated for 1, 2, 3, 4, 6, and 9 hours at room temperature. Overall, samples collected in the AM period showed greater richness and alpha-diversity compared to samples collected in the PM period. While a significant effect of time was detected in all hindgut regions, the effect increased from cecum to distal colon. When using two vendor-origin microbiomes as a biological variable, its effect size vastly outweighed the effect size of the time samples spent at room temperature, the time of day samples were collected, and the position within the colon from which samples were collected.

**Conclusions:** This study has highlighted multiple scenarios encountered in microbiome research that may affect outcome measures of microbial diversity and composition. Unexpectedly, delayed time to sample cold storage up to nine hours did not affect the alpha or global beta diversity of fecal sample. We then presented evidence of location- and time-dependent effects within the hindgut on microbial richness, diversity, and composition. We finally demonstrated a relatively low effect size of these technical factors when compared to a primary experimental factor with large intergroup variability.

## Background

The gut microbiome (GM) is a collection of resident microorganisms that inhabit the gastrointestinal tract of a host organism [1]. In health, the GM confers numerous benefits to the host, including diversification of dietary compounds, transformation of xenobiotics, colonization resistance against pathogens, and many more. Research has also shown that the GM can influence many pathophysiological and disease processes within the host including obesity [2], inflammatory bowel disease [3], colon cancer [4], mental health disorders [5], and autism [6], potentially alleviating or exacerbating disease processes. Due to ethical and practical concerns surrounding the use of humans and other larger mammals in microbiome studies, the mouse has become an essential research model to unravel and understand how the GM can influence host health and disease. A common experimental component in this area of research is the collection of fecal biomass for molecular analysis of bacterial composition, often through sequencing 16S rRNA amplicon libraries. Fecal boluses represent a noninvasive, easily acquired, and highly informative sample, enabling high-density, longitudinal studies, and data generation and analysis have become relatively standardized.

While murine fecal collection is pivotal for GM studies, there are a number of practical factors that must be considered in order to ensure sound scientific data when performing fecal collection. Some GM studies include large numbers of mice in order to achieve high statistical power (reduced type II error), potentially increasing the time required to collect freshly evacuated fecal boluses from each mouse. As some mice may require a long period of time to defecate, this increases the time that the first collections may sit at room temperature while the remaining fecal samples are collected. Similarly, logistical factors or simple oversight may also result in fecal samples experiencing increased time at room temperature before being appropriately stored. Studies examining the stability of bacterial communities of equine fecal samples demonstrated changes in beta diversity after six hours at room temperature [7,8]. Another study examining the long-term effects of temperature on the microbial composition in dog feces demonstrated significant changes in alpha diversity and microbial relative abundance after two weeks [9]. Studies looking at human samples have concluded that bacterial communities in feces remain stable up to 24 hours at both room temperature and 4°C [10,11]. Another group compared the microbial communities of pig fecal samples collected directly from the rectum and stored in liquid nitrogen and samples stored at room temperature for 3 hours and found no difference in microbial ecological indices [12]. Surprisingly, the effect of increased time at room temperature on the relative abundance of bacteria within murine samples remains unreported.

Samples are also frequently collected at necropsy during terminal procedures, often from the rectum or descending (distal) colon as this represents the colonic contents closest to becoming a freshly evacuated fecal bolus. However, a fecal bolus may not be present in the desired region of the colon requiring that the sample be taken from a different region such as the middle or proximal colon. While it has been demonstrated that GM bacterial composition can follow a diurnal pattern within cecum [13] and feces [14,15], it remains unknown if this pattern is conserved across other colonic regions. Similarly, a researcher may lose control of the time at which a terminal sample is collected. Many researchers are aware of the diurnal rhythms present within the host and microbiome, and control for this by performing terminal procedures and sample collection at a uniform time of day. Such procedures in mice are frequently performed in the morning, as anecdotal evidence suggests that more feces will be present in the colon due to nighttime feeding behavior compared with fecal collection in the evening. IACUC protocols and study guidelines may also require that mice be euthanized due to reaching a humane endpoint. This may require euthanasia in the evening while other samples will be collected in the morning, potentially confounding fecal data from this animal when compared to the rest of the cohort. Thus, numerous scenarios exist wherein circumstances dictate that sample collection varies in terms of hindgut location or time of day.

Differences in the GM of mice from various rodent suppliers have been documented and characterized by us and others [12,13]. These differences in alpha diversity, beta diversity, and bacterial composition of the GM have also been shown to influence multiple research and disease models [16–18]. Thus, we used the difference between two supplier-origin microbiomes as a standard biological variable with a large effect size, against which to compare other experimental variables related to sample collection. Additionally, this provides an assessment of the reproducibility of any detected effects of duration at room temperature, time of day, or location in the hindgut across multiple specific pathogen-free (SPF) microbiomes. As such, the use of two different supplier-origin SPF microbiomes in parallel experiments enhances rigor and provides validation of findings common to both GMs, which would suggest broad applicability.

## Methods

### Ethics Statement

This study was conducted in accordance with the recommendations set forth by the Guide for the Care and Use of Laboratory Animals and was approved by the University of Missouri institutional Animal Care and Use Committee (MU IACUC protocol 36781).

### Mice

C57BL/6J mice were purchased from Jackson Laboratory (Bar Harbor, ME, USA) and set up as breeding trios. The pups from these trios were cross-fostered within 24 hours of birth onto CD-1 dams either harboring GM_Low_ or GM_High_ to transfer their respective GMs to the surrogate pups. The surrogate pups generated from these cross-fostered litters were confirmed to have been colonized with the GM of their respective CD-1 donor surrogate dam via 16S rRNA gene amplicon sequencing and were used as colony founders. The mice used in this study were the 5^th^ generation of these colonies. Briefly, the colonies of CD-1 mice that were used as GM donors (i.e., surrogate dams for cross-fostered C57BL/6J mice) were originally generated by implanting CD-1 embryos into pseudopregnant C57BL/6J (Jackson Laboratory, Bar Harbor, ME, USA), or C57BL/6NHsd (Envigo, Indianapolis, IN, USA) surrogate dams [19]. CD-1 pups from these embryo transfers acquired the supplier-origin microbiome of their respective dams and served as founders for outbred colonies that have been maintained under barrier conditions at the MU Mutant Mouse Resource and Research Center (MMRRC, Columbia, MO, USA) for several years. To be clear, GM_Low_ and GM_High_ represent the microbiome originally acquired from C57BL/6J (Jackson) and C57BL/6NHsd (Envigo) mice, respectively.

All C57BL/6J mice used in this study were housed two animals per cage under barrier conditions in microisolator cages (Thoren, Hazleton, PA, USA) with aspen chip bedding. Mice had *ad libitum* access to irradiated LabDiet 5053 maintenance feed (LabDiet, St. Louis, MO, USA), and autoclaved tap water. The facility maintains all animals under 12:12 light/dark cycle. Mice were found to be free of *Bordetella bronchiseptica; Filobacterium rodentium; Citrobacter rodentium; Clostridium piliforme Corynebacterium bovis*; *Corynebacterium kutscheri*; *Helicobacter* spp.; *Mycoplasma* spp.; *Pasteurella pneumotropica*; *Pneumocystis carinii*; *Salmonella* spp.; *Streptobacillus moniliformis*; *Streptococcus pneumoniae*; adventitious viruses including H1, Hantaan, KRV, LCMV, MAD1, MNV, PVM, RCV/SDAV, REO3, RMV, RPV, RTV, and Sendai viruses; intestinal protozoa including *Spironucleus muris*, *Giardia muris*, *Entamoeba muris*, trichomonads, and other large intestinal flagellates and amoebae; intestinal parasites including pinworms and tapeworms; and external parasites including all species of lice and mites via quarterly sentinel testing.

### Sample Collection

#### Room temperature study

At 50 days of age, mice were individually placed into clean cages and allowed to naturally defecate for a period of time not exceeding 15 minutes. Single pellets were retrieved and placed into individual microcentrifuge tubes using autoclaved wooden toothpicks. One fecal pellet from each cage was immediately snap frozen in liquid nitrogen representing time 0. The remaining pellets were placed at room temperature (21°C) for 1, 2, 3, 6, and 9 hours before being snap-frozen in liquid nitrogen at the appropriate time. After snap-freezing, samples were stored at -80°C until processing.

#### Colon position study

Mice were euthanized via CO_2_ asphyxiation followed by cervical dislocation. The lower gastrointestinal tract (cecum to anus) was collected, trimmed of mesenteric fat, and photographed. Coli were photographed, and colon lengths and number of fecal pellets within each colon were collected. Individual fecal pellets from the proximal, mid, and distal colon and cecal contents were collected for 16S rRNA sequencing.

#### DNA extraction

DNA was extracted using a modified QIAamp PowerFecal Pro DNA extraction kits (QIAGEN). Samples were collected into 2.0 mL round-bottom microcentrifuge tubes with a single 0.5 cm steel ball. Samples were homogenized at for 10 min at 30 Hz using a TissueLyser II (QIAGEN). DNA extraction continued per manufacturer instructions. DNA yields were quantified via fluorometry (Qubit 2.0, Invitrogen, Carlsbad, CA) using quant-iT BR dsDNA reagent kits (Invitrogen). When appropriate, DNA yields were normalized to 3.51 ng/μL using Buffer C6 (QIAGEN)

#### 16S rRNA library preparation and sequencing

Library preparation and sequencing were performed at the University of Missouri Genomics Technology Core. Bacterial 16S rRNA amplicons were constructed via amplification of the V4 region of the 16S rRNA gene with universal primers (U515F/806R) [20] previously developed against the V4 region, flanked by Illumina standard adapter sequences. PCR was performed as 50 µL reactions containing 100 ng metagenomic DNA, dual-indexed forward and reverse primers (0.2 µM each), dNTPs (200 µM each), and Phusion high-fidelity DNA polymerase (1U, Thermo Fisher). Amplification parameters were 98°C^(3^ ^min)^ + [98°C^(15^ ^sec)^ + 50°C^(30^ ^sec)^ + 72°C^(30^ ^sec)^] × 25 cycles + 72°C^(7^ ^min)^ [21]. Amplicon pools were combined then purified by addition of Axygen Axyprep MagPCR clean-up beads to an equal volume of 50 µL of amplicons and incubated for 15 minutes at room temperature. Products were washed multiple times with 80% ethanol and the pellet was resuspended in 32.5 µL EB buffer (Qiagen), incubated for two minutes at room temperature, and then placed on the magnetic stand for five minutes. The final amplicon pool was evaluated using an Advanced Analytical Fragment Analyzer automated electrophoresis system, quantified using quant-iT HS dsDNA reagent kits, and diluted according to the Illumina standard protocol for sequencing as 2×250 bp paired-end reads on the MiSeq instrument.

#### Informatics

All 16S rRNA amplicons were processed using the Quantitative Insights Into Microbial Ecology 2 (QIIME2) framework v2021.8 [22]. Illumina adapters and primers were trimmed from forward and reverse reads with cutadapt [23]. Untrimmed sequences were discarded. The paired-end reads were then truncated to 150 base pairs and denoised into amplicon sequence variants (ASVs) using DADA2 [24]. Paired-end reads were merged based on a minimum overlap of 12 base pairs. Merged sequences were filtered to between 249 and 257 base pairs in length. Unique sequences were then assigned a taxonomic classification using a sklearn algorithm and the QIIME2-provided 99% non-redundant SILVA v138 [25] reference database trimmed to the 515F/806R [20] region of the 16S rRNA gene. The resulting feature table of ASV counts per sample was rarefied to a uniform depth of 10,734 features per sample.

#### Microbiome Analysis

All microbiome and statistical analyses were performed using the described libraries within R v4.2.2 [26]. All code can be accessed at https://github.com/ericsson-lab/fecal_collection_study. Univariate data were reported as mean ± standard error (SE). Data were first assessed for normality using the Shapiro-Wilkes test. Single-factor analyses were performed using T tests if data were normally distributed and Wilcoxon-Rank Sum tests if not. When testing for differences in data with two or more factors, the appropriate multifactorial analysis of variance (ANOVA) was used. Significant differences were further explored using *post hoc* Tukey HSD testing.

#### Alpha Diversity Metrics

Alpha diversity metrics (Chao-1 and Shannon Indices) were calculated using the *microbiome* [27] and *vegan* [28,29] libraries, respectively.

#### Beta Diversity

A distance matrix using weighted distances was generated using the *vegdist* function (*vegan*) from a quarter-root transformed feature table. Multivariate (compositional) data was assessed for differences between groups using a permutational analysis of variance (PERMANOVA) with 9,999 permutations. Principal coordinate analysis (PCoA) was performed using the *ape* library [30] with a Calliez correction.

### Differential abundance

#### Room temperature study

Differences in taxonomic composition within each GM were determined using two concurrent differential abundance (DA) tools, serial ANOVA and ANCOM-BC2. The approach to test for differentially abundant taxa within each GM differed between the two vignettes. Serial ANOVA testing (*rstatix* [31]) was performed using the following model within each taxa: ‘anova_test(rel_abund ∼ time_point)’. BH-corrected *p* values < 0.05 were considered significant. ANCOM-BC2 testing (*ancombc* [32,33]) was performed using the fixed effect ‘time_point’ and group variable ‘time_point’. Significant differentially abundant taxa were determined based on a BH-corrected *p* value (*q*) < 0.05. Structural zeroes were identified based on presence/absence in relation to the group variable.

#### Colon position study

Serial ANOVA testing (*rstatix* [31]) was performed using the following model within each taxa: ‘anova_test(rel_abund ∼ time_point * sample_type)’. BH-corrected *p* values < 0.05 were considered significant. *Post hoc* Tukey HSD tests were performed using the same model. A *p* < 0.05 was considered significant.

## Results

### C57BL/6J mice colonized with two standardized complex GMs differ in richness, diversity, and composition

To determine the effects of room temperature incubation and spatiotemporal sample collection on microbial community analysis, we utilized C57BL/6J (B6) mice colonized with one of two standardized complex microbiomes maintained by the NIH Mutant Mouse Resource & Research Center at the University of Missouri. Relative to each other, these GMs exhibit large differences in community richness (i.e., number of unique amplicon sequence variants [ASVs]), diversity, and composition. B6 mice colonized with GM_High_ exhibited greater community richness (*p* < 0.001) and Shannon diversity (*p* < 0.001) relative to GM_Low_ (**Figure 1A-B**). Differences in community composition using weighted (Bray-Curtis) distances were also observed (F = 39.35, *p* < 0.001) and visualized using principal coordinate analysis (PCoA, **Figure 1C**.). Modest sex-dependent effects on community richness were observed (**Figure S1A**), but not diversity or community composition (**Figure S1B-C**). To determine whether the sex-dependent effect on community richness, which is determined using ASV counts, corresponded with the taxonomic composition of each sex, we identified the shared genera between males and females of GM_Low_ and GM_High_. GM_Low_ and GM_High_ mice shared 88.31% (68/77) and 93.02% (80/86), respectively, of genera between males and females (**Figure S1D-E**). Shared genera in GM_Low_ displayed an average prevalence of 78.6% (median = 100%) and average abundance of 1.47% (median = 0.19%, **Figure S1D-E**). In GM_High_, shared genera displayed an average prevalence of 76.5% (median = 100%) and average abundance of 1.25% (median = 0.23%, **Figure S1D-E**). Given the high proportion of shared taxa and relatively low effect size of sex between males and females in both GMs, sex was removed as a factor in the remainder of the study.

**Figure 1.**
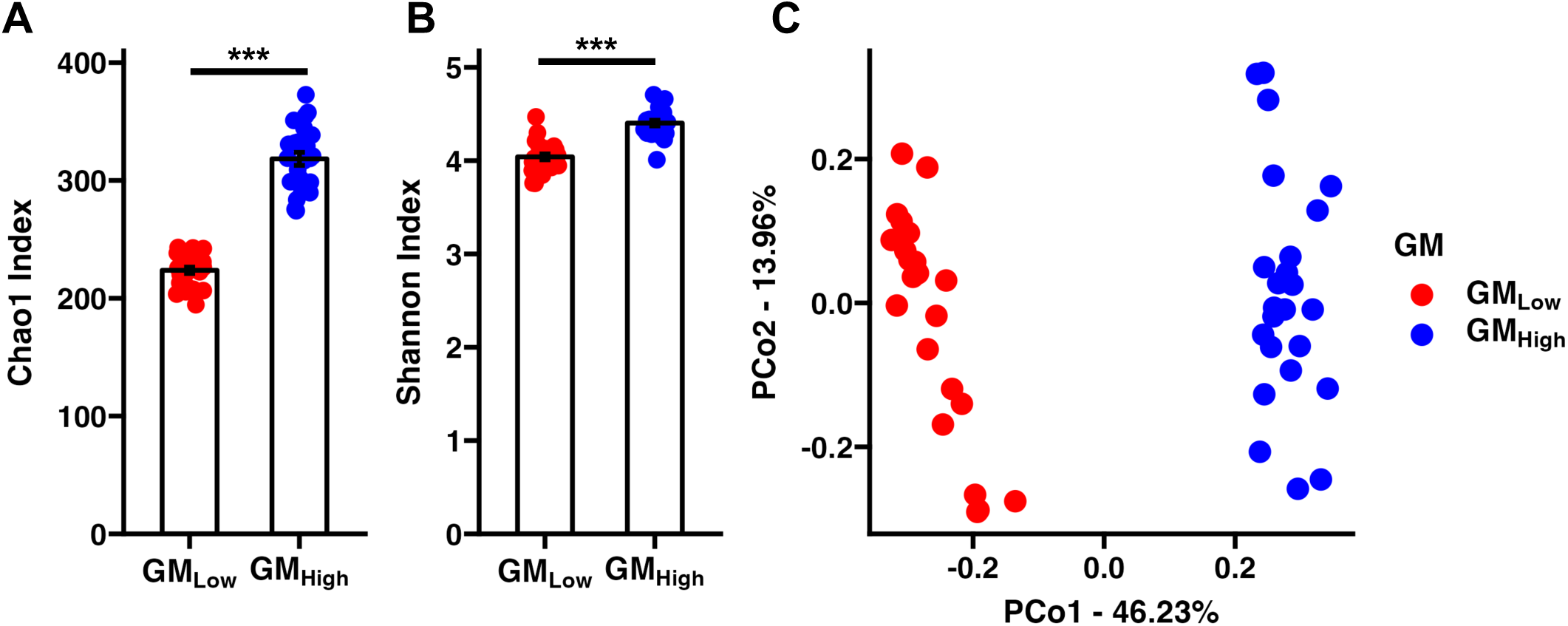
Standardized complex GMs differ in alpha and beta diversity. (A) Dot plot depicting significant GM-dependent differences in Chao1 richness. *p* < 0.001. Wilcoxon Rank-Sum test. (B) Dot plot depicting significant GM-dependent differences in Shannon diversity. *p* < 0.001. T test. (B) Principal coordinate analysis depicting significant GM-dependent differences in community composition using Bray-Curtis distances. F = 39.01 *p* < 0.001. One-way PERMANOVA. *n* = 24-25 mice/GM.

### Delayed freezing up to nine hours does not affect alpha or global beta diversity or taxonomic composition of the fecal microbiome

First emulating the real-world scenario in which fecal samples are not immediately frozen after collection, fecal pellets were collected from pair-housed mice and immediately (0 hr) snap-frozen in liquid nitrogen or stored at room temperature (∼21°C) for a period of 1, 2, 3, 6, or 9 hours before freezing. The maximum time spent at room temperature (9 hr) was selected to mimic the length of a standard workday. We then assessed whether prolonged storage of mouse fecal samples at room temperature affects common 16S rRNA microbial community analysis outcome measures. Longitudinal analysis of intra-cage alpha diversity metrics revealed a significant effect of GM but not time spent at room temperature on community richness (**Figure 2A**; GM: F = 350.74, *p* < 0.001; Timepoint: F = 0.66, *p* = 0.418) and Shannon diversity (**Figure 2B**; GM: F = 174.25, *p* < 0.001; Timepoint: F = 2.15, *p* = 0.144) suggesting that storage at room temperature for extended period of time does not affect alpha diversity.

**Figure 2.**
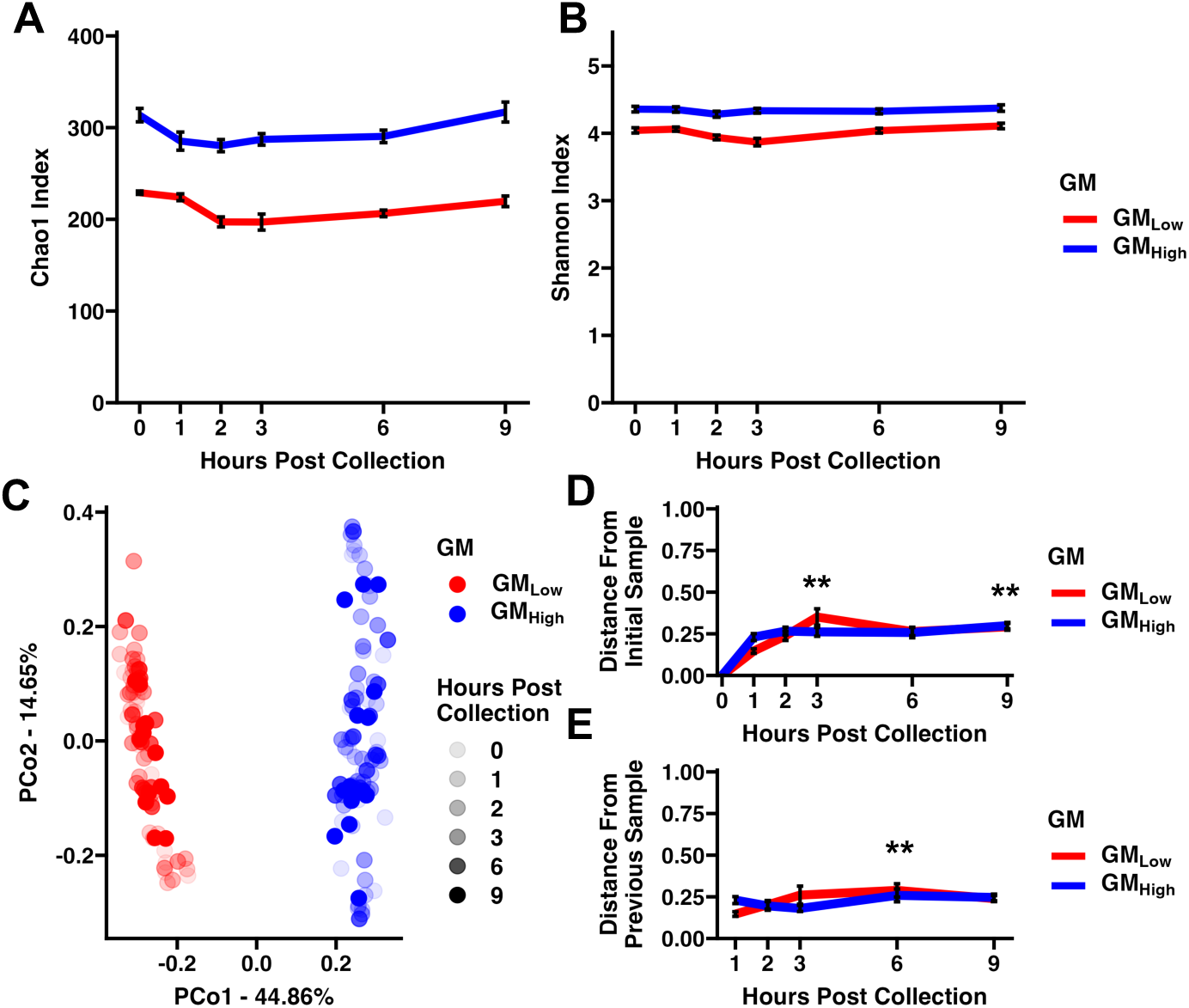
Room-temperature incubation of murine fecal samples affects beta but not alpha diversity. **(A)** Line plot depicting intracage Chao1 richness across time. GM: F = 350.74, *p* < 0.001. Timepoint: F = 0.66, *p* = 0.418. Two-way ANOVA. **(B)** Line plot depicting intracage Shannon diversity across time. GM: F = 174.3, *p* < 0.001. Timepoint: F = 2.15, *p* = 0.144. Two-way ANOVA. (C) Principal coordinate analysis depicting between sample diversity across time using Bray-Curtis distances. GM: F = 126.7, *p* < 0.001. Timepoint: F = 1.27, *p* = 0.170. Two-way PERMANOVA. **(D)** Line plot depicting intracage Bray Curtis dissimilarity from initial sample (T0). GM: F = 0.07, *p* = 0.791. Time point: F = 7.18, *p* = 0.008. ** *p* < 0.01 relative to 1 hr, Tukey post hoc test. **(E)** Line plot depicting intracage Bray Curtis dissimilarity from previous timepoint. GM: 0.07, *p* = 0.792. Time point: F = 6.218, *p* = 0.014. ** *p* < 0.01 relative to 1 hr, Tukey post hoc test. *n* = 13-14 cages/GM.

Next assessing beta diversity, we identified GM-but not time-dependent differences in overall community composition using weighted distances (**Figure 2C**; GM: F = 126.65, *p* < 0.001; Timepoint: F = 1.27, *p* = 0.170). This demonstrates that the length of time a fecal sample incubates at room temperature does not affect global beta diversity. Even when stratifying by GM, no time-dependent effects on beta diversity were observed (**Figure S2A-B**). We then explored beta diversity at a more granular level by determining the intra-cage beta diversity across each timepoint using Bray-Curtis distances. GM_Low_ samples frozen at 3 hours post-collection visually appeared to have increased dissimilarity to all other time points (**Figure S2C**), whereas GMHigh samples frozen 9 hours post-collection visually appeared to have an increased dissimilarity to all other time points (**Figure S2D**). To make practical comparisons, we next assessed Bray-Curtis dissimilarity of intra-cage samples relative to the initial collection (0 hr, **Figure 2D**). We found that distance from time 0 was significantly affected by time spent at room temperature (F = 3.19, *p* = 0.03), but not GM (F = 1.25, *p* = 0.266). *Post hoc* Tukey tests revealed significant differences in distance from baseline (0 hr) between hours 1 and 3 (*p* = 0.009) and hours 1 and 9 (*p* = 0.022). The gradual increase in dissimilarity from hour 0 prompted us to assess dissimilarity from the previous timepoint to identify the timepoint(s) at which the largest changes in community composition occur (**Figure 2E**). A significant effect of time on distance from previous time point was observed (GM: F = 0.07, *p* = 0.792; Timepoint: F = 6.22, *p* = 0.014), however, upon *post hoc* comparison, only one significant comparison was observed. These data collectively suggest that no large shifts in composition but rather gradual increases in dissimilarity from immediate freezing may occur.

Lastly, we evaluated whether changes in taxonomic relative abundance occurred during extended storage at room temperature. Visual inspection of the average phylum-level relative abundance of both GM_Low_ and GM_High_ across timepoints indicated shifts in the abundance of *Bacteroidota* and *Bacillota* between hours 1-3 (**Figure 3A-B**). Using analysis of composition of microbiomes with bias correction 2 (ANCOM-BC), we found that, at the phylum level, no resolved phyla were differentially abundant across timepoints in either GM. ANCOM-BC2 also identifies structural zeros – taxa present in at last one group and absent in at least one group. Only one, unresolved bacterial phyla was identified as a structural zero in both GM_Low_ and GM_High_ (**Supplementary File 1**). We then performed the same analysis at the family level. Again, visual inspection of the family abundance across timepoints suggested that, as at the phylum level, the abundance of major families like *Lachnospiraceae* (phylum *Bacillota*) and *Muribaculaceae* (phylum *Bacteroidota*) changed between 1-3 hours post-collection (**Figure 4C-D**), however, ANCOM-BC2 determined that no families were differentially abundant across timepoints in either GM. Within GM_Low_ and GM_High_, 21 and 18 families, respectively, were identified as structural zeroes being present in at least one timepoint but absent in another. Of the dominant families (average relative abundance > 1%) present in GM_Low_ and GM_High_, only *Erysipelotrichaceae* (phylum *Bacillota*) was found to be a structural zero within GM_High_ (**Supplementary File 1**). These data were corroborated with serial ANOVAs within GM_Low_ and GM_High_. Only one taxa (*Anaerovoracaceae*) significantly differed across timepoints (Benjamini-Hochberg [BH]-corrected *p* < 0.05, **Supplementary File 2**). Collectively, these data support that fecal sample storage at room temperature for up to 9 hours does not affect sample richness or diversity. Subtle effects on beta diversity were observed, however, storage at room temperature did not affect the taxonomic abundance at the phylum or family levels.

**Figure 3.**
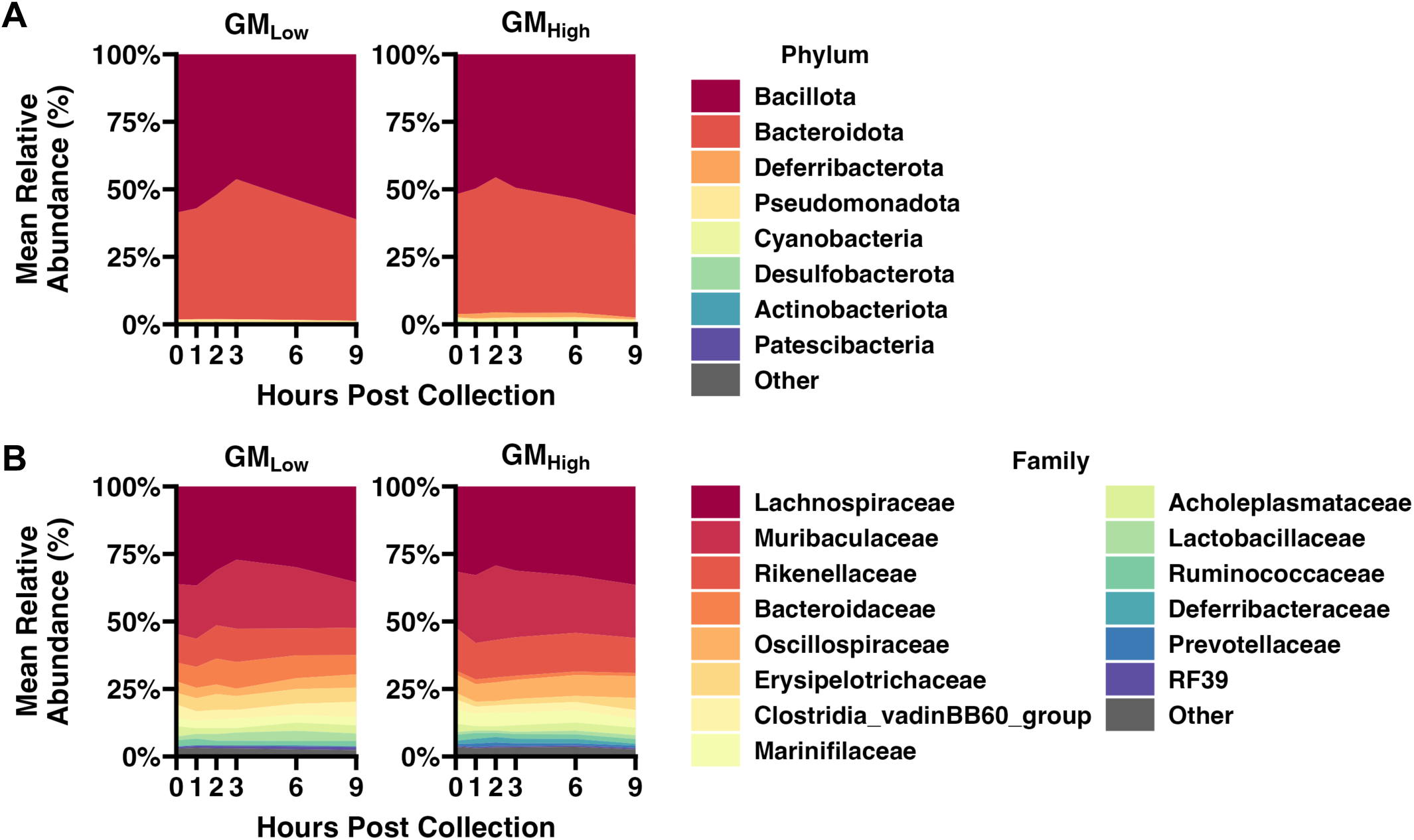
Delayed freezing does not affect taxonomic relative abundance. **(A)** Area plot depicting mean relative phylum abundance of the dominant taxa (> 0.1%) in GM_Low_ and GM_High_ across time points. **(B)** Area plot depicting mean relative family abundance of the dominant taxa (> 1%) in GM_Low_ and GM_High_ across time points. *n* = 13-14 cages/GM

**Figure 4.**
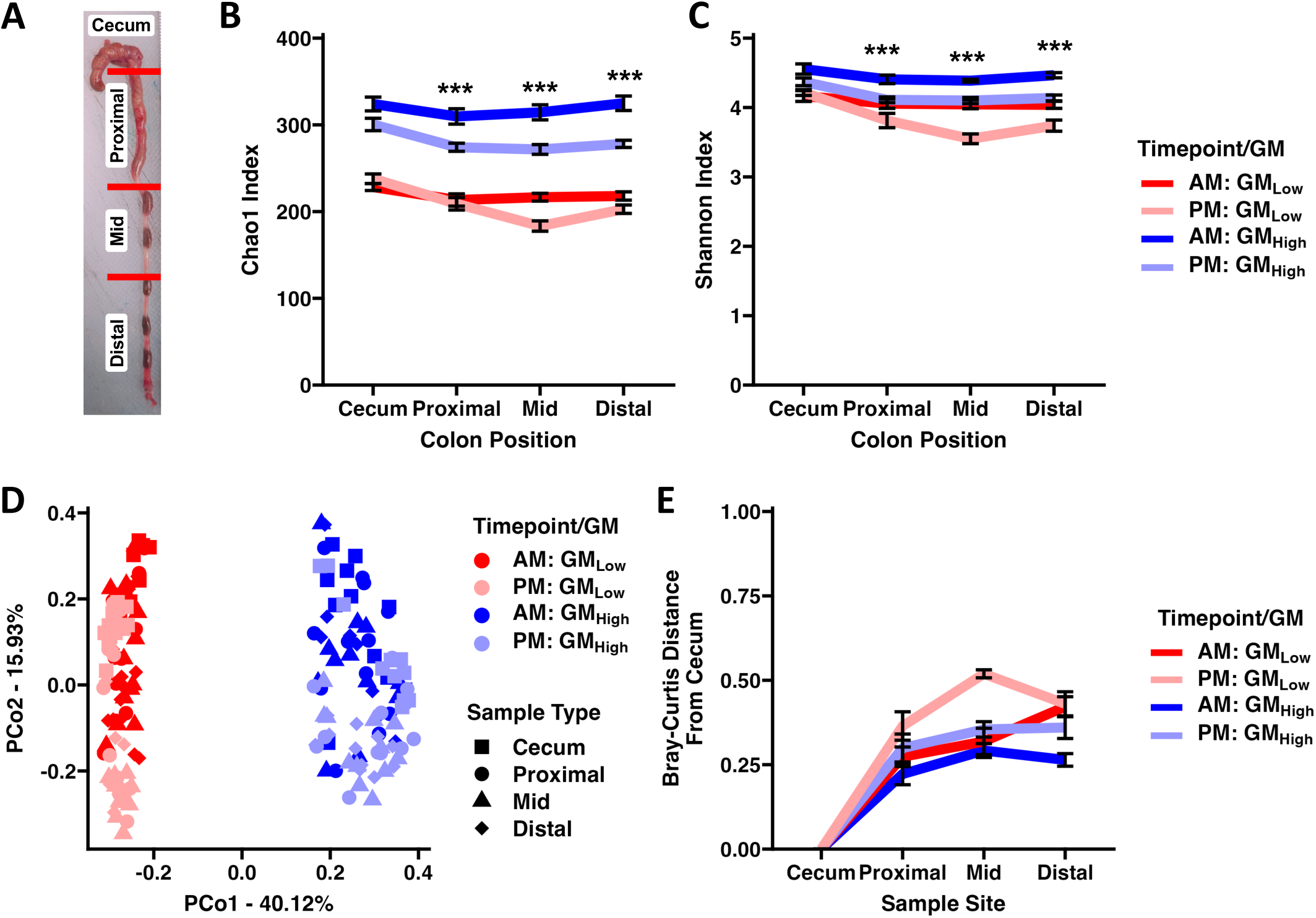
Terminal sample position and collection period affect alpha and beta diversity. **(A)** Representative image of terminal hindgut collection depicting where the indicated samples were collected for the cecum and proximal, mid, and distal colon. *\*\*\*p* < 0.001 relative to cecum, Tukey *post hoc* test. **(B)** Line plot depicting intrasubject Chao1 richness across sample location. GM: F = 738.29, *p* < 0.001. Time point: F = 57.23, *p* < 0.001. Sample type: F = 14.01, *p <* 0.001. Three-way ANOVA. *\*\*\*p* < 0.001 Tukey *post hoc* test. **(C)** Line plot depicting intrasubject Shannon diversity across sample location. GM: F = 138.3, *p* < 0.001. Timepoint: F = 71.88, *p* < 0.001. Sample type: F = 19.45, *p <* 0.001. Three-way ANOVA. **(D)** Principal coordinate analysis depicting between sample diversity across sample types and time using Bray-Curtis distances. GM: F = 162.47, *p* < 0.001. Timepoint: F = 26.32, *p* < 0.001. Sample type: F = 8.63, *p <* 0.001. One-way PERMANOVA. **(E)** Line plot depicting intrasubject Bray Curtis dissimilarity from cecum. GM: F = 24.21, *p* < 0.001. Timepoint: F = 23.29, *p* < 0.001. Sample type: F = 8.68, *p <* 0.001. Three-way ANOVA. *n* = 10-12 mice/time point/GM

### Spatiotemporal differences in the microbial ecology of the murine hindgut

We next assessed spatiotemporal effects by collecting cecal contents and colonic contents from the proximal, mid, and distal colon of GM_Low_ and GM_High_ mice at 07:00 (AM) and 16:00 (PM) (**Figure 4A**). With regard to the total number of distinct fecal boluses present in the colon, no significant main effects of GM or collection period were detected, although there was a significant interaction between GM and collection period on the number of fecal boluses present (GM × collection period: F = 11.50, *p* = 0.001). Specifically, coli from GM_Low_ mice contained more fecal boluses than coli from GM_High_ mice when collected in the AM (*p* = 0.005, **Figure S3**).

A longitudinal analysis of community richness across sample locations revealed significant effects of GM (F = 738.3, *p* < 0.001), sample location (F =14.01, *p* < 0.001), and time of collection (F = 57.22, *p* < 0.001) (**Figure 4B**). Significant interactions of GM × sample type (F = 17.40, *p* < 0.001) and sample type × collection period (F = 4.66, *p* = 0.004) were also observed. As expected, GM_High_ samples were richer than those from GM_Low_ across all sample sites. Consistent with the known diurnal rhythmicity of the gut microbiome [13,14], AM samples exhibited greater richness relative to PM samples in both GMs. When comparing sample locations, cecal samples were richer than proximal (*p* < 0.001), mid (*p* < 0.001), and distal colon samples (*p* < 0.001). Shannon diversity also differed between GMs (F = 136.3, *p* < 0.001), sample locations (F = 19.45, *p* < 0.001), and collection period (F = 71.88, *p* < 0.001) (**Figure 4C**). Similar to community richness, GM_High_ samples displayed greater diversity than GM_Low_. Samples collected in the AM were more diverse than those collected in the PM. When comparing sample locations, cecal samples exhibited greater diversity relative to proximal, mid, and distal colon samples. When comparing Shannon diversity, a significant interaction of sample location and collection period was also observed (F = 5.02, *p* = 0.002). *Post hoc* Tukey tests revealed fifteen significant interactions which, of note, included collection period-dependent effects on proximal (*p* < 0.001), mid (*p* < 0.001), and distal (*p* = 0.002) colon samples.

Using a three-way PERMANOVA, we identified significant GM-(F = 162.5, *p* < 0.001), sample location- (F = 8.63, *p* < 0.001), and collection period-dependent effects (F = 26.32, *p* < 0.001) on global community composition (**Figure 4D**). Significant GM × sample location (F = 2.40, *p* = 0.005) and GM × collection period (F = 12.14, *p <* 0.001) interactions were also observed. These sample location- and collection period-dependent effects of community composition were also observed when individually assessing beta diversity within GM_Low_ and GM_High_ (**Figure S4A-B**).

Focusing our investigation next on intrasubject beta diversity, we determined the Bray-Curtis dissimilarity of proximal, mid, and distal colon samples relative to cecal samples (**Figure 4E**). We identified GM- (F = 24.21, *p* < 0.001), collection period- (F = 23.39, *p* < 0.001), and sample location-dependent (F = 8.68, *p* < 0.001) effects on the Bray-Curtis dissimilarity from cecal samples. Compared to proximal colon, samples collected from the mid (*p* < 0.001) and distal (*p* = 0.002) colon exhibited greater dissimilarity from the cecum. Comparing the intrasubject Bray-Curtis distances between sample locations revealed greater dissimilarity in samples collected in the PM (GM_Low_: 0.281 ± 0.047; GM_High_: 0.211 ± 0.035) than those collected in the AM (GM_Low_: 0.229 ± 0.037; GM_High_: 0.177 ± 0.028) within both GMs (**Figure S3C-D**). While collection period-dependent effects on community composition were observed in every sample location of both GMs, the distance between community centroids was lowest in cecal samples relative to more distal samples (**Figure S3E-F**).

We then assessed collection period-dependent effects on taxonomic abundance across sample sites using serial two-factor ANOVA testing. At the phylum level (**Figure S4**), *Actinobacteria* and *Patescibacteria* significantly differed between collection periods, and p*ost hoc* analysis revealed that within the distal colon, only *Actinobacteria* differed between collection periods. Four phyla including *Bacteroidota*, *Bacillota*, *Desulfobacterota*, and *Pseudomonadota* significantly differed between sample locations (**Figure S4**). The relative abundance of fourteen (29.2%) and nineteen families (38.8%) families differed between collection period and sample location, respectively, in GM_Low_ (**Figure 5**). Seven families including *Bifidobacteriaceae*, *Erysipelotrichaceae*, *Lachnospiraceae*, *Muribaculaceae*, *Peptococcaceae*, *RF39*, and *Rikenellaceae* exhibited collection period-dependent effects in at least one sample site. Of those families, only *RF39* exhibited collection period-dependent effects on relative abundance in the cecum. The relative abundance of *Actinobacteria*, *Patescibacteria*, *Bacillota*, *Bacteroidota*, *Pseudomonadota*, *Cyanobacteria*, and *Defferibacterota* differed between collection periods, whereas, only *Bacillota*, *Bacteroidota*, *Pseudomonadota*, and *Desulfobacterota* differed between sample locations in GM_High_ (**Figure S4**). Collection period-dependent effects on the relative abundance of both *Bacteroidota* and *Bacillota* were observed in the proximal, mid, and distal colon but not in the cecum. Only *Desulfobacterota* exhibited collection period-dependent effects on relative abundance in the cecum. The abundance of nineteen (32.8%) and fourteen (24.1%) families significantly differed between collection periods and sample locations (**Figure 5**). Of those families, only *Anaerovoracaceae* and *Desulfovibrionaceae* differed between collection periods in the cecum (**Figure 5**). Many differences between sample locations were observed in GM_Low_ and GM_High_. A comprehensive list of taxa differing between sample location and collection period has been provided in **Supplementary File 3.**

**Figure 5.**
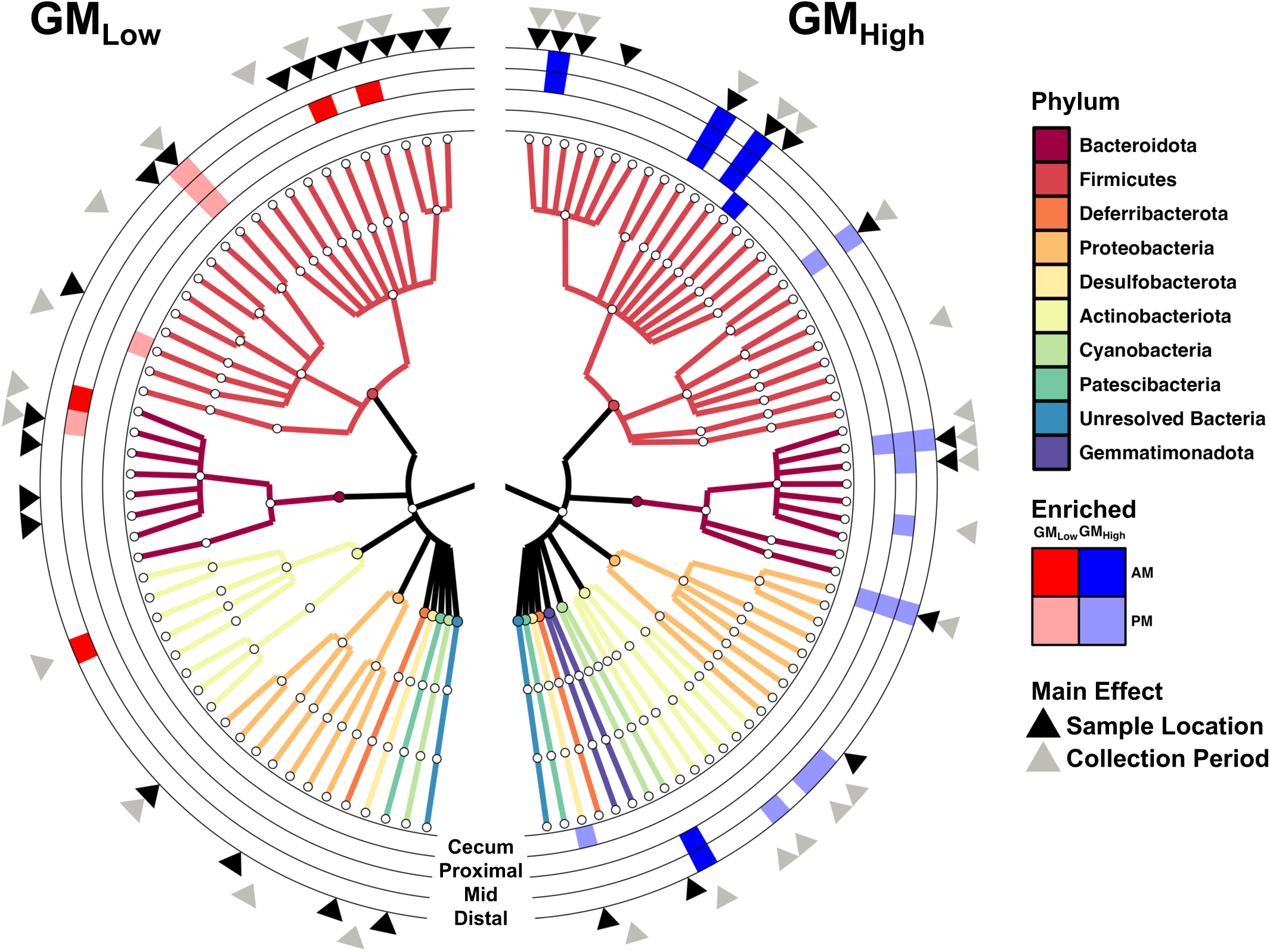
Family level abundance differs between sample location and collection period. Family-level phylogenetic tree for GM_Low_ (*left*) and GM_High_ (*right*). Phylum-level classification denoted by branch color. Rings depict collection period-dependent effect on family level abundance in the indicated sample location. Black and gray arrows indicate overall sample location- and collection period-dependent effects on family relative abundance. *n* = 10-12 mice/time point/GM.

### Primary experimental factor contributes greatest intergroup variability

Finally, we characterized the magnitude of statistical effect size attributable to these sample collection and handling factors in the context of our primary experimental factor (GM). To compare effect sizes, we calculated the partial eta squared (η_p_^2^: small = 0.01, medium = 0.06, large > 0.14 [34]) for the appropriate main effects from three common microbiome outcome measures: Chao-1 Index, Shannon Index, and a PERMANOVA (Bray-Curtis). In the room temperature experiment, GM contributed an average effect size of 0.580 across the three tests while time left at room temp contributed a four-fold less average effect size of 0.109 (**Figure 6A**). In our spatiotemporal analysis of lower GI samples, the collection period and sample location contributed an average effect size of 0.208 and 0.177, respectively, whereas GM contributed an average effect size of 0.556 (**Figure 6B**). These data indicate that while variables like time spent at room temperature, sample location, and collection period do contribute moderate effects on common microbiome outcome measures, in the context of an experimental group with high intergroup variability, these effects are muted.

**Figure 6.**
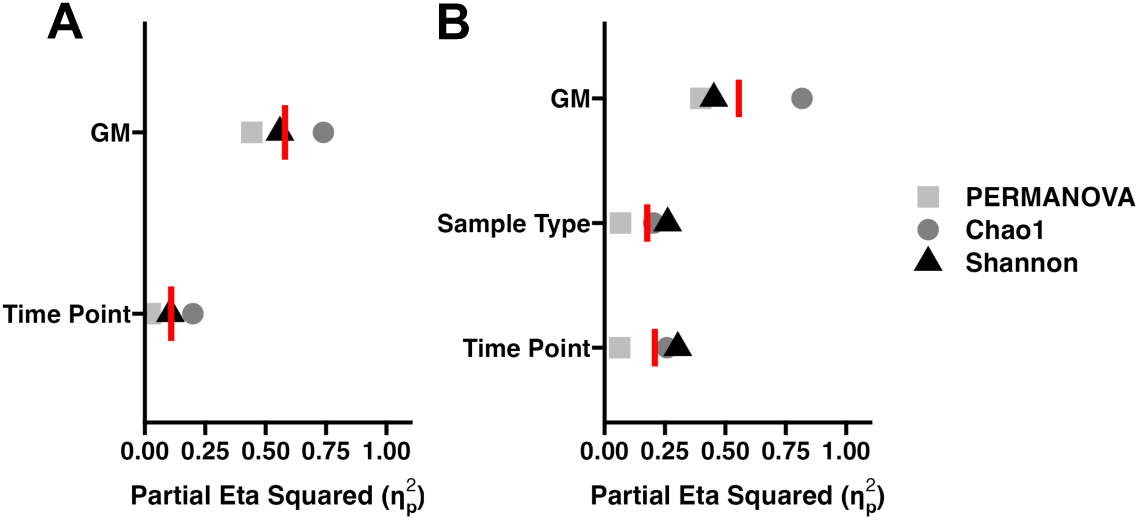
Primary experimental factor contributes high effect size relative to technical factors **(A)** Dot plot depicting the effect size (η_p_^2^) of GM and time point to microbial richness, diversity, and composition. Red line depicts the average η_p_^2^ value. **(B)** Dot plot depicting the effect size (η_p_^2^) of GM, sample type, and time point to microbial richness, diversity, and composition. Red line depicts the average η_p_^2^ value.

## Discussion

Standardization of sample collection and handling methods will improve the rigor and reproducibility of microbiome science. Here we leveraged a robust model comprising C57BL/6J mice colonized with one of two standardized complex GMs to determine whether practical scenarios often encountered in microbiome studies meaningfully affect outcome measures of microbial ecology. With respect to one another, GM_Low_ and GM_High_, exhibit robust differences in alpha **(Figure 1A-B**) and beta diversity (**Figure 1C**) [19,21]. We sought to identify whether the time a fecal sample spends at room temperature, sample location within the murine hindgut, or time of day terminal collections are performed affect the expected differences between two distinct microbial communities. Our data reveal that while these technical variables affect microbiome outcome measures, the magnitude of these effects are modest relative to the effect of treatment groups with high intergroup variability.

In a laboratory setting, there are many situations that may prevent the immediate cold storage of a murine fecal sample including long *in vivo* fecal collection sessions, transport between facilities, or general human error (e.g., samples left on benchtop). Previous investigations have identified host species-dependent effects on the microbial ecology of fecal samples stored at ambient temperature for extended periods of time. Equine fecal samples exhibit decreased alpha diversity and increased community dissimilarity relative to fresh samples after six hours due in part to rapid proliferation of *Enterococcaceae* and *Bacillaceae* [7,8]. Similar analyses of human fecal samples demonstrate that short-term storage at room temperature does not affect sample alpha or between-group beta diversity, however, this stability does not persist when samples are stored for longer than 24 hours [10,11]. Our data demonstrate that storing a mouse fecal sample at ambient temperature for up to nine hours does not affect intra-cage alpha diversity (**Figure 2A-B**) while minimally affecting community composition (**Figure 2C-E**). We observed increased community dissimilarity in microbial composition relative to samples that were frozen immediately across time (**Figure 2D**), however, this variation is likely no greater than the expected intrasubject variation observed upon repeat sampling [35]. This stability may be due to the relatively small size of murine fecal boluses allowing for rapid water loss. Future investigations may seek to characterize the stability of the murine fecal microbiome left at room temperature beyond nine hours, however, given the ease and accessibility of collecting these samples in the laboratory setting, investigators should collect fresh samples and immediately snap-freeze to minimize potential shifts in microbial composition.

A rigorous approach to microbiome science should include collecting fecal boluses from the same region of the hindgut to minimize spatial difference in microbial composition. While consistently collecting samples from the same region is ideal, it is not always feasible. If for example, no sample is present in the hindgut region of interest upon necropsy, investigators may instead collect a sample from an immediately adjacent region. While groups have characterized the microbial diversity of the upper and lower murine GI tract [36,37], few have provided a granular assessment of hindgut biogeography [38]. Our data demonstrate that in two standardized complex GMs, the hindgut position from which a sample is collected affects community richness (**Figure 4B**), diversity (**Figure 4C**), and composition (**Figure 4D**, **Figure 5**). For example, in both GM_Low_ and GM_High_ relative *Lachnospiraceae* abundance peaked in the cecum and generally decreased towards the distal colon whereas *Muribaculaceae* relative abundance was low in cecal samples but increased distally. Given the present data, one must consider how spatial differences within the hindgut microbiome affect the outcome measures of interest when considering alternative samples.

Diurnal oscillations of microbial diversity and taxonomic composition in the gut pose an additional factor to consider in microbiome science. While maintaining consistency in the time of day at which terminal samples are collected is ideal, some experimental protocols may prohibit this. Take for example an investigator that elects to perform all terminal collections in the morning (i.e., late dark/early light phase). If an animal reaches a humane endpoint (e.g., weight loss, tumor size, moribund) in the afternoon (i.e., late light/early dark phase), the terminal samples from the animal euthanized in the afternoon would not be in the same stage of microbial periodicity as samples collected in the morning. Consistent with previous reports [13,14], our data demonstrate that samples collected in the morning exhibit increased richness and diversity relative to those collected in the afternoon (**Figure 4B-C**). Considering these diurnal fluctuations, investigators may consider collecting samples from the region exhibiting the least amount of temporal variation. We propose that the cecal microbiome provides the least amount of variability between collection periods. Cecal samples did not differ in either community richness or diversity between collection periods. While significant differences in community composition were observed between timepoints within individual sample locations (**Figure S4E-F**), AM and PM cecal samples clustered closer to one another within both GMs relative to all other sample sites (**Figure 4D**). Furthermore, when assessing intrasubject beta diversity relative to cecal samples (**Figure 4E**), samples collected in the afternoon displayed greater dissimilarity relative to those collected in the morning. These data collectively suggest that when posed with an experimental situation in which asynchronous terminal sample collection is necessary, cecal samples provide the least amount of temporal variability in microbial diversity and composition. However, if animals are not expected to reach a human endpoint during the study, consistency in both time of day and hindgut region of interest are recommended for terminal sample collections.

Here we have leveraged a model of population-level variability of the gut microbiome [19,21] to determine whether the technical challenges of microbiome research presented in this study affect known differences in intergroup variability of microbial diversity and composition. In doing so, we compared relative effect sizes across multiple microbiome outcome measures using partial eta squared. We found that our primary experimental variable (GM) contributed the largest proportion of overall variance compared to factors associated with sample collection and handling (**Figure 6**). While these technical factors contributed a moderate (η_p_^2^ > 0.06) to large (η_p_^2^ > 0.14) effect size, their contribution was considerably smaller relative to the primary experimental factor. The degree to which these technical factors affect microbiome outcomes likely depends on the expected degree of variability between primary treatment groups. Our data suggest that if the anticipated variance between primary treatment groups is high, the contribution of the sample collection and handling factors to outcome variability is expected to be less.

## Conclusion

In this study, we have modeled three scenarios encountered in microbiome research that may affect outcome measures of microbial diversity and composition. We show that delayed freezing of murine fecal samples for up to nine hours does not affect alpha diversity, global beta diversity, or taxonomic composition. We then provided a granular assessment of differences in spatiotemporal microbial composition within the hindgut. Our data revealed sample location- and time-dependent effects on microbial richness, diversity, and taxonomic composition. Finally, we demonstrated that the effect size of these technical factors is low relative to a primary experimental factor with known large intergroup variability. Collectively, these data are of great value to the field as they contribute to the ongoing effort to improve rigor and reproducibility in microbiome research and provide guidance in the event of unforeseen circumstances related to sample collection.

## Data Availability

16S rRNA sequencing data are available at the Sequence Read Archive under the BioProject number PRJNA980714. All code can be accessed at https://github.com/ericsson-lab/fecal_collection_study.

## Acknowledgements

The authors would like to thank the MU MMRRC (NIH U42 OD010918) for generously providing the CD-1 mice.

## Author Contributions

Conceptualization – AE

Data Collection – AE, KG, ZM, AR, BG, GT

Data Analysis – AE, KG, ZM

Writing manuscript – AE, KG, ZM

Reviewed manuscript - AE, KG, ZM

**Supplementary Figure 1.** Modest sex-dependent differences in community richness between GM_Low_ and GM_High_. (A) Dot plot depicting significant GM-and dependent differences in Chao1 richness. GM: F = 266.63, *p* < 0.001. Sex: F = 5.97, *p* = 0.011. \*\*\**p* < 0.001 Two-way ANOVA. (B) Dot plot depicting significant GM-dependent differences in Shannon diversity. GM: F = 67.29, *p* < 0.001. Sex: F = 0.78, *p* = 0.382. \*\*\**p* < 0.001 Two-way ANOVA. (C) Principal coordinate analysis depicting significant GM-dependent differences in community composition using Bray-Curtis distances. GM: F = 40.84, *p* < 0.001. Sex: 2.13, *p* = 0.071. Two-way PERMANOVA. Venn diagram depicting shared and sex-specific genera detected in GM_Low_ (D) and GM_High_ (E). (F) Dot plot depicting the prevalence and abundance of the core and sex-specific genera detected in GM_Low_ and GM_High_. *n =* 11-13 mice/sex/GM

**Supplementary Figure 2.** Delayed freezing does not affect beta diversity. (A) Principal coordinate analysis depicting between sample diversity across time using Bray-Curtis distances within GM_Low_. Timepoint: F = 1.78, *p* = 0.104. One-way PERMANOVA. (B) Principal coordinate analysis depicting between sample diversity across time using Bray-Curtis distances within GM_High_. Timepoint: F = 1.56, *p* = 0.127. One-way PERMANOVA. Heatmap depicting average intracage Bray-Curtis dissimilarity between all timepoints in GM_Low_ (C) and GM_High_ (D). *n* = 13-14 cages/GM.

**Supplementary Figure 3.** Mice colonized with low richness microbiome contain more fecal boluses in the morning relative to mice colonized with a high richness microbiome. Dot plot depicting the number of fecal boluses observed in the hindgut of animal sacrificed in the morning and afternoon necropsies. GM: F = 2.693, *p* = 0.108. Time point: F = 0.108, *p* = 0.744. GM×Time point: F = 11.50, *p* = 0.001. Two-way ANOVA. ** *p* < 0.01. *Post hoc* Tukey test. *n* = 10-12 mice/time point/GM.

**Supplementary Figure 4.** Spatiotemporal differences in murine hindgut microbial beta diversity. **(A)** Principal coordinate analysis depicting between sample diversity across sample locations and collection periods using Bray-Curtis distances within GM_Low_. Sample type: F = 9.03, *p* < 0.001. Time point: F = 25.24, *p* < 0.001. Two-way PERMANOVA. **(B)** Principal coordinate analysis depicting between sample diversity across sample locations and collection periods using Bray-Curtis distances within GM_High_. Sample type: F = 2.51, *p* = 0.002. Time point: F = 9.88, *p* < 0.001. Two-way PERMANOVA. **(C)** Heatmap depicting average intrasubject Bray Curtis dissimilarity between sample types collected in the AM (*top left*) and PM (*bottom right*) in GM_Low_. (D) Heatmap depicting average intrasubject Bray Curtis dissimilarity between sample types collected in the AM (*top left*) and PM (*bottom right*) in GM_High_. **(E)** Principal coordinate analyses of each sample type depicting collection period-dependent differences in beta diversity within GM_Low_. Ballot box symbols represents the centroid of PCo1 and PCo2. The centroid distance (CD) between timepoints along all coordinates is indicated. One-way PERMANOVA F and *p* values are depicted below each plot. **(F)** Principal coordinate analyses of each sample type depicting collection period-dependent differences in beta diversity within GM_High_. Ballot box symbols represents the centroid of PCo1 and PCo2. The centroid distance (CD) between timepoints along all coordinates is indicated. One-way PERMANOVA F and *p* values are depicted below each plot. *n =* 10-12 mice/time point/GM.

**Supplementary Figure 5.** Phylum level abundance differs between sample location and collection period. Phylum-level phylogenetic tree for GM_Low_ (*left*) and GM_High_ (*right*). Phylum-level classification denoted by branch color. Rings depict collection period-dependent effect on phylum level abundance in the indicated sample location. Black and gray arrows indicate overall sample location- and collection period-dependent effects on family abundance. *n* = 10-12 mice/time point/GM.

## Abbreviations

*16SrRNA*: 16S ribosomal RNA
*ASV*: Amplicon Sequence Variants
*DA*: Differential Abundance
*GI*: Gastrointestinal Tract
*GM*: Gut Microbiome
*IACUC*: Insitutional Animal Care and Use Committee
*PCoA*: Principal Coordinate Analysis
*SPF*: Specific Pathogen-Free
*QIIME 2*: Quantitative Insights Into Microbial Ecology 2

