## Supplementary figures and images for "Effect size of delayed freezing, diurnal variation, and hindgut location on the mouse fecal microbiome relative to a standardized biological variable"

### Figure S1

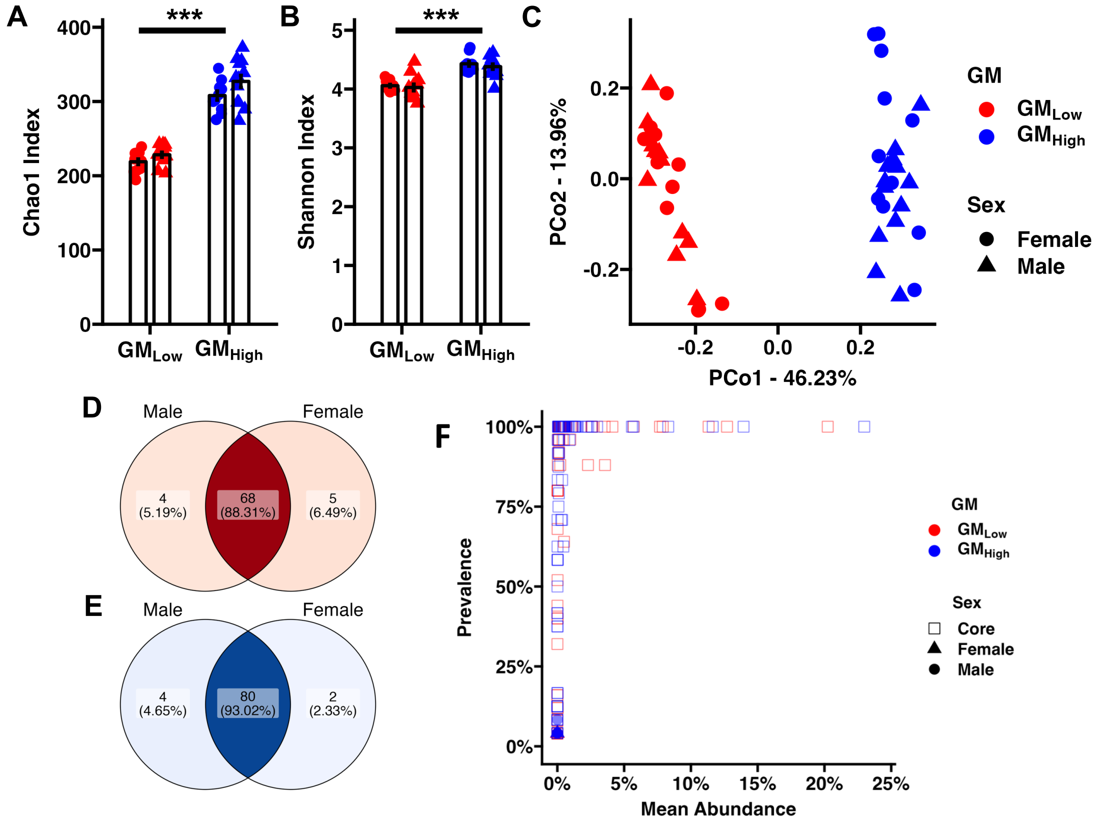

### Figure S2

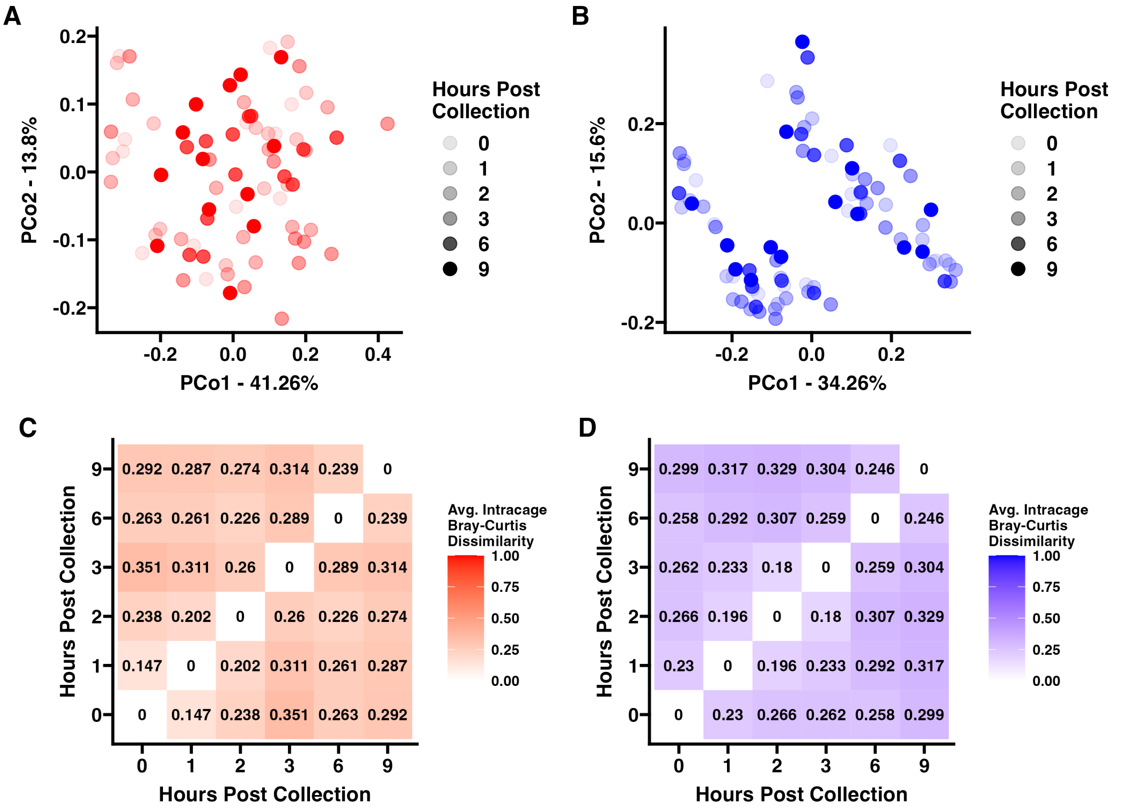

### Figure S3

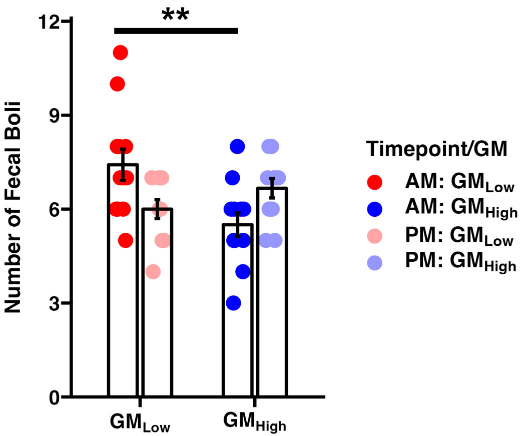

### Figure S4

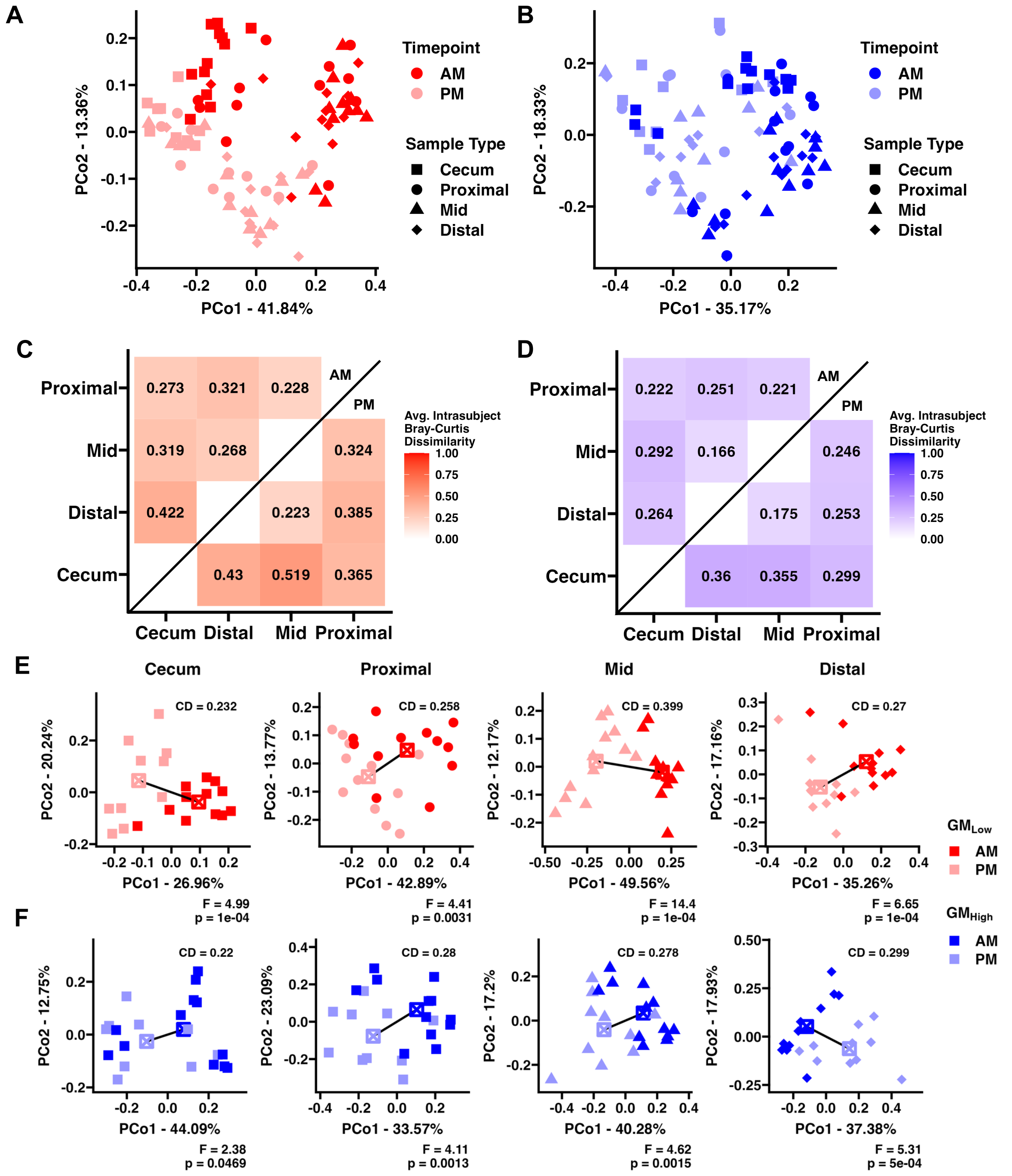

### Figure S5

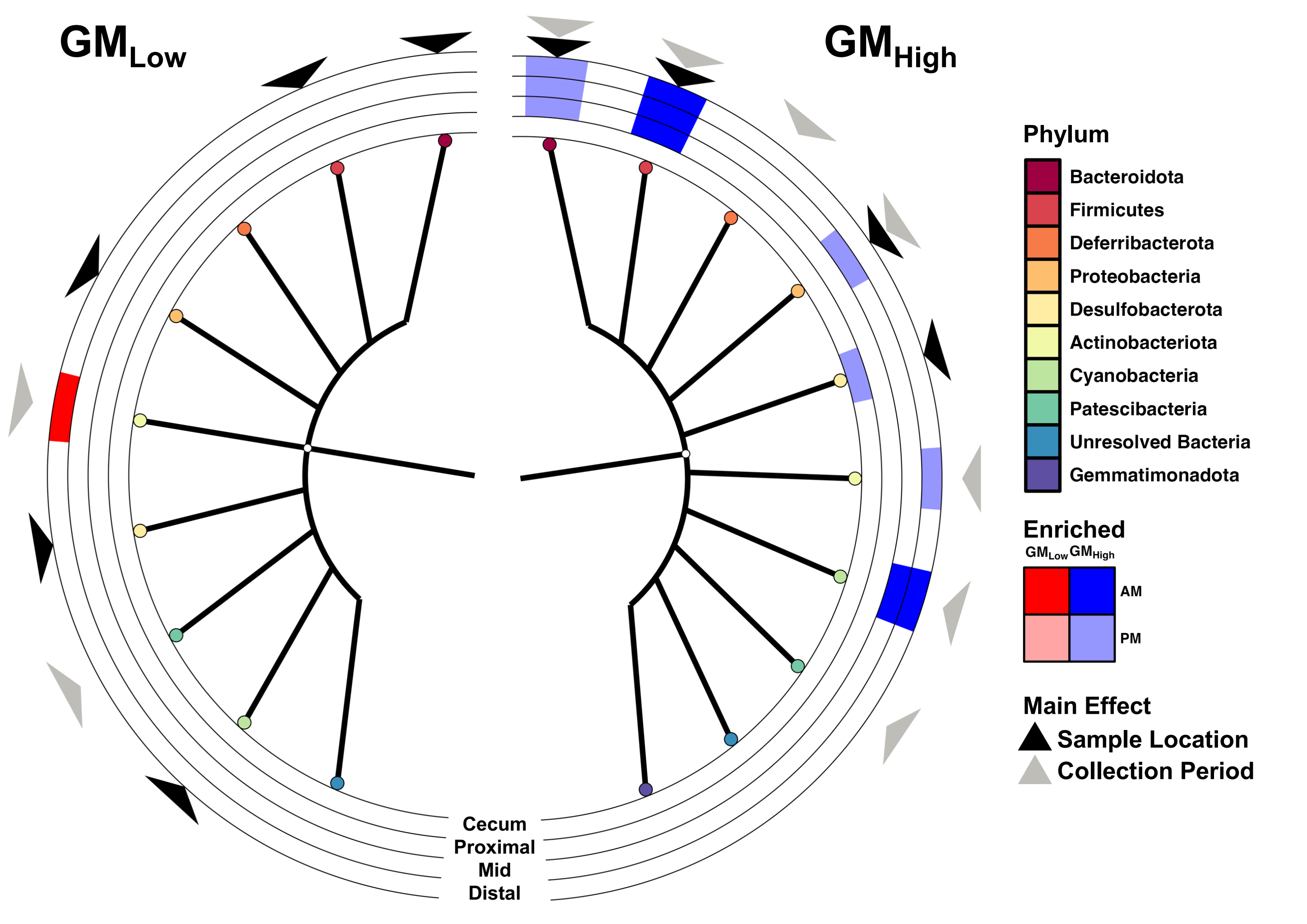
